# DNA replication stress stratifies prognosis and enables exploitable therapeutic vulnerabilities of HBV-associated hepatocellular carcinoma: an *in silico* strategy towards precision oncology

**DOI:** 10.1101/2021.09.04.458962

**Authors:** Xiaofan Lu, Jialin Meng, Yujie Zhou, Haitao Wang, Xinjia Ruan, Yi Chen, Yuqing Ye, Liwen Su, Xiaole Fan, Hangyu Yan, Liyun Jiang, Fangrong Yan

**Affiliations:** State Key Laboratory of Natural Medicines, Research Center of Biostatistics and Computational Pharmacy, China Pharmaceutical University, Nanjing, P.R. China; Department of Urology, The First Affiliated Hospital of Anhui Medical University; Institute of Urology & Anhui Province Key Laboratory of Genitourinary Diseases, Anhui Medical University, Hefei, Anhui, P.R. China; Division of Gastroenterology and Hepatology, Key Laboratory of Gastroenterology and Hepatology, Ministry of Health, Renji Hospital, School of Medicine, Shanghai Jiao Tong University, Shanghai Institute of Digestive Disease, Shanghai, P.R. China; Cancer Center, Faculty of Health Sciences, Center for Precision Medicine Research and Training, University of Macau, Macau SAR, P.R. China; Department of Radiology, The Second Affiliated Hospital of Nantong University, Nantong, P.R. China; Department of Biostatistics, The University of Texas MD Anderson Cancer Center, Texas, USA

**Keywords:** Hepatitis B virus, hepatocellular carcinoma, DNA replication stress, prognostic stratification, precision oncology

## Abstract

**Background:** Hepatitis B virus (HBV), the main risk factor for hepatocellular carcinoma (HCC) development, integrates into the host genome, causing genetic instability, which may trigger malignancies to exhibit chronic DNA replication stress, providing exploitable therapeutic vulnerabilities. Therefore, customizing prognostication approach and expanding therapeutic options are of great clinical significance to HBV-associated HCCs.

**Methods:** A robust machine-learning framework was designed to develop a DNA replication stress-related prognostic index (*PI*_*RS*_) based on 606 retrospectively collected HBV-associated HCC cases. Molecular profiles and drug response of HCC cell lines were leveraged to predict therapeutic targets and agents for patients with high mortality risk.

**Results:** Compared with established population-based predictors, *PI*_*RS*_ manifested superior performance for prognostic prediction in HBV-associated HCCs. Lower *PI*_*RS*_ tightly associated with higher expression of HBV oncoproteins, activated immune/metabolism pathways and higher likelihood of responding to immunotherapy; while higher *PI*_*RS*_ showed co-occurrence manner with elevated *Ki-67* progression marker and cancer stemness, and significantly enriched in DNA replication stress, cell cycle pathways, chromatin remodeling regulons, and presented an ‘immune-cold’ phenotype with unfavorable clinical outcome. Through large-scale *in silico* drug screening, four potential therapeutic targets (*TOP2A, PRMT1, CSNK1D*, and *PPIH*) and five agents including three topoisomerase inhibitors (teniposide, doxorubicin, and epirubicin) and two *CDK* inhibitors (JNJ-7706621 and PHA-793887) were identified for patients with high *PI*_*RS*_.

**Conclusions:** Overall, *PI*_*RS*_ holds potential to improve the population-based therapeutic strategies in HCC and sheds new insight to the clinical management for those HBV carriers; current analytic framework provides a roadmap for the rational clinical development of personalized treatment.

## INTRODUCTION

Hepatocellular carcinoma (HCC) ranks as the fourth leading cause of cancer-related deaths, and hepatitis B virus (HBV) and hepatitis C virus (HCV) infection are the major risk factors for HCC development ^1^. Unlike HCV, an RNA virus which never integrates into the host genome, HBV is a small DNA virus who frequently integrates into the genome of the host hepatic cells and progressively contributes to hepatocarcinogenesis. However, the precise mechanism by which it causes HCC remains unknown and the optimal therapeutic regimens for treating HBV-associated HCC have not yet been established ^2, 3^. Most importantly, HBV carriers have a worse prognosis, with relative risks of 6.27 and 2.2 for mortality of HCC and chronic liver disease, respectively ^4, 5^. Nevertheless, most prognostic predictors were developed based on the entire HCC population without exploring tailored clinical management for patients with higher risk ^6-9^, which is insufficient for precise prognostic stratifications and medication. Therefore, an urgent need is pursued to tailor effective management for HBV-associated HCC.

Integrated HBV DNA in the host genome triggers genotoxicity and genome instability, leading to selective advantages for tumour progression ^10^. Of note, genome stability is carefully surveilled by DNA damage response (DDR), inducing DNA repair and cell-cycle checkpoints that stall the replication of damaged cells, and often causes replication stress, which in turn exacerbates genome instability and therefore potentiates oncogenic transformation ^11^. There is a growing compendium of novel therapeutics that target DDR and cell cycles ^12, 13^, and Dreyer *et al*. recently reported that high replication stress offered therapeutic opportunities for pancreatic cancer ^14^. Nevertheless, evidence is scarce regarding how to exploit DNA replication stress for HCC, especially for those HBV carriers who were considered to have higher sensitive to DNA damage ^15^.

To address above issues, for the first time to our best knowledge, we developed a robust DNA replication stress-related prognostic index (*PI*_*RS*_) for HBV-associated HCC and predicted potential therapeutic targets as well as agents for those patients with high mortality risk. Specifically, *PI*_*RS*_ exhibited superior predictive performance in 606 cases across four independent cohorts comparing to previously established population-based signatures. Furthermore, four potential therapeutic targets (*i*.*e*., *TOP2A, PRMT1, CSNK1D*, and *PPIH*) and five agents including three topoisomerase inhibitors (*i*.*e*., teniposide, doxorubicin, and epirubicin) and two *CDK* inhibitors (*i*.*e*., JNJ-7706621 and PHA-793887) were identified for patients with high *PI*_*RS*_, which holds potential to improve the population-based therapeutic strategies in HCC and sheds new insight to the clinical management for those HBV carriers.

## MATERIALS AND METHODS

### RNA-sequencing cohort

Molecular profiles of patients diagnosed with HCC were collected from the Cancer Genome Atlas under project of TCGA-LIHC. The raw paired-end reads FASTQ files were acquired for transcriptome quantification and virus detection. DNA methylation matrix was obtained from XENA archive (https://xenabrowser.net/). Segment of copy number was downloaded from FireBrowse (http://firebrowse.org/). Somatic mutations, clinicopathological characteristics, and clinical outcome were retrieved from cBioPortal (https://www.cbioportal.org/). A total of 359 primary tumors and 50 adjacent normal samples shared with multi-omics profiles and survival information were identified. Another RNA-Seq cohort, CN-LIHC, includes 318 paired tumor and normal liver tissues from 159 Chinese HCC patients infected by HBV were also retrieved from the literature ^16^; transcriptome expression measured as the number of fragments per kilobase million (FPKM), somatic mutation, gene-level proteomic data and clinical outcome were collected. Additionally, the liver cancer-RIKEN (LIRI-JP) project from the International Cancer Genome Consortium (ICGC) portal (https://dcc.icgc.org/) with 216 HCCs was downloaded for clinical information, transcriptome FPKM value and somatic mutations; Hepatitis B antigen (HBAg) status for each donor was retrieved from the literature ^17^, leading to 62 HBV-associated HCCs for LIRI-JP cohort.

### RNA analysis

#### Data preprocessing

The raw paired-end reads were aligned to GRCh37/hg19 human reference genome using MOSAIK. The mapped reads in genomic features were counted using HTSeq package and annotated in GENCODE27 to generate transcriptome raw counts. We chose the “union” mode of HTSeq so as to overcome the non-strand-specific RNA sequencing issue in kit used by TCGA. We then converted raw counts to FPKM and further into transcripts per kilobase million (TPM) values, signal of which were more similar to those quantified from microarray and more comparable between cases ^18^.

#### Virus detection

We leveraged VirusSeq to align the RNA-Seq libraries to both human and HBV genomes to detect HBV and quantify viral oncoprotein expression ^19^. At least one of four oncoproteins, *i*.*e*., HBVgp3-X, HBVgp2-S, HBVgp2-pre-S1/S2, and HBVgp4-c was identified in 103 out of 359 HCCs in TCGA-LIHC cohort. According to our previous study ^20^, we established a new and comprehensive variable to explain the original viral expression by principal component analysis (PCA). Specifically, we developed *HBV*_*pca*_ which considered the first and second PCs that explained 80.1% and 15.3% of the variation, respectively. Mathematically, let *E*_*ij*_ represents log_2_(FPKM + 1) of oncoprotein *j* in sample *i*, and *C*_*jk*_ denotes the corresponding coefficient of oncoprotein (*HBV*_*j*_, *j* ∈ {1,2,3,4}) for principal component *k* (*k* ∈ {1,2}). A matrix ℳ was calculated as below:

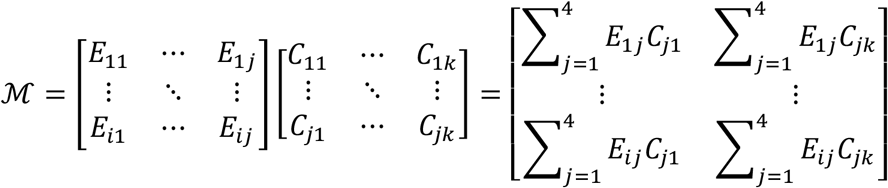

where *i* = 1, …, 103; *j* = 1,2,3,4; *k* = 1,2. We took the row sum of ℳ and derived the *HBV*_*pca*_ as the following:

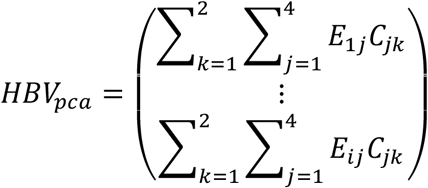

### Microarray cohorts

We retrospectively collected the gene expression profiles of frozen tumor tissue samples of HCC with available HBV infection status from two microarray cohorts, including GSE14520 and GSE121248. Specifically, 212 out of 221 HCC samples from GSE14520 showed active viral replication chronic carrier or chronic carrier of HBV, and all 70 cases in GSE121248 carried chronic HBV. The OS information for GSE14520 cohort was retrieved from the literature ^21^.

### Batch effect removal

The potential cross-cohort batch effect was removed by *removeBatchEffect()* function using R package “*limma*” ^22^, and PCA was harnessed to evaluate whether batch effect was removed appropriately (**Figure S1a-b**).

### Cancer cell lines

Expression profile, somatic mutation, and description of human cancer cell lines (CCLs) were downloaded from the dependency map (DepMap) portal (https://depmap.org/portal/) ^23^. The CERES scores for 17,645 genes in 990 CCLs were acquired from DepMap (CRISPR [Avana] Public 21Q2). CERES computationally estimated gene-dependency from CRISPR–Cas9 essentiality screens while adjusting the copy number-specific effect ^24^. Drug response of CCLs were obtained from PRISM Repurposing Primary Screen (19Q4) which archived in DepMap. The primary PRISM Repurposing dataset records pooled-cell line chemical-perturbation viability screens for 4,518 compounds against 578 CCLs; the secondary dataset records pooled-cell line chemical-perturbation viability screens for 1,448 compounds screened against 489 CCLs undergoing eight dose points: 0.00061, 0.00244, 0.00977, 0.0391, 0.15625, 0.625, 2.5, and 10uM. The sensitivity to specific compound was measured as log_2_FoldChange at replicate level for data relative by negative-control wells (dimethyl sulfoxide, DMSO) from the same CCL on the same detection plate; replicate collapsed log_2_FoldChange values relative to DMSO were corrected for experimental confounders using ComBat. Additionally, dose-response curves were fit to the secondary replicate-level viability data with the measurement of the area under the dose response curve (AUC); lower log_2_FoldChange or AUC indicate increased sensitivity to treatment. Generally, we filtered out 24 CCLs with primary HCC (one HBs-antigen carrier [ACH-000475]), namely, HCC cell lines (HCCLs). After sample match according to different analytic purpose, we obtained 22 HCCLs shared with expression profile, 20 HCCLs shared with CERES scores, 19 HCCLs shared with primary drug sensitivity, 14 HCCLs shared with secondary dose-finding, and 17 HCCLs shared with dose response curves.

### Development of a prognostic replication stress-related signature

We retrieved 21 replication stress signatures from the literature ^14^. To develop robust prognostic signature in HBV-associated HCC, we preliminary screened prognostic replication stress-related genes in TCGA-LIHC training cohort via univariate Cox regression (*P*<0.01) and a bootstrap approach was leveraged to test the robustness of prognostic value. Specifically, 80% of samples were randomly extracted without replacement from the entire cohort and univariate analysis was performed to these subsets. Such bootstrap process was repeated 1,000 times and genes that were incorporated in more than 800 times of the resampling were kept. We then applied random survival forest (RSF) by using R package “*randomForestSRC*” to further narrow down the prognostic gene list. In this analysis, 1,000 trees were built using a log-rank score splitting algorithm and features were selected by variable hunting procedure with variable importance. The RSF was independently repeated 1,000 times and genes combination with largest concordance index (C-index) were considered as optimal signature ^25^. Consequently, *PI*_*RS*_ was then calculated individually via a linear combination of the expression of selected genes, weighted by the corresponding Cox regression derived coefficients. Mathematically, let *e*_*ij*_ represents the expression of specific replication-stress related gene *i* in sample *j*, and *β*_*i*_ represents the regression coefficient for gene *i*; let *X* = (*x*_1_, …, *x*_*j*_)^T^ denotes the original *PI*_*RS*_ as the following:

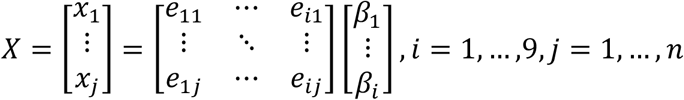

Original *PI*_*RS*_ was further min-max normalized and times 10 to ensure comparability among different cohorts as the following:

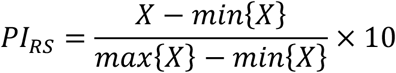

### Existing prognostic signatures and classifications for comparison

We retrospectively collected four published HCC prognostic signatures for comparison, including Li *et al*.’s three-gene signature ^6^, Yan *et al*.’s four-gene signature ^7^, Hu *et al*.’s five-gene signature ^8^, and Chen *et al*.’s nine-gene-pair signature ^9^; genes in Hu *et al*.’s signature failed to be fully matched in LIRI-JP or GSE14520 cohort. To assess the efficiency and robustness of *PI*_*RS*_ in survival prediction, we randomly resampled 80% of the cases from TCGA-LIHC, CN-LIHC, LIRI-JP and GSE14520 cohorts for 10,000 times; prognostic indexes were calculating by multiplying gene expression values through their corresponding reported coefficients and summing these values, and the overall C-index was further calculated for comparison. Previously published molecular classifications of HCC, including Boyault’s classification ^26^, Chiang’s classification ^27^, Hoshida’s classification ^28^, and Désert’s classification ^29^, were also predicted through nearest template prediction by using R package “*CMScaller*” ^30^.

### Bioinformatics analyses

Differential expression analyses were conducted using R package “*limma”* ^*22*^, and a meta-analysis using random effect model (REM) was harnessed to obtain a comprehensive landscape of expression pattern across different cohorts ^31^.

To gain biological understanding of *PI*_*RS*_, we conducted transcriptome-based pathway analysis (Pathifier) and proteome-based gene set enrichment analysis (GSEA) with background of hallmark gene sets (h.all.v7.4.symbols, https://www.gsea-msigdb.org/gsea/msigdb/) ^32^. Specifically, Pathifier was performed using R package “*pathifier*” with input of expression profiles from HBV-associated tumors and adjacent normal samples in TCGA-LIHC and CN-LIHC cohorts, and pathway deregulation score (PDS) which could exhibit the degree of deregulation of certain biological process was calculated. GSEA was performed on CN-LIHC cohort through R package “*clusterProfiler*” ^33^ (*P*<0.05 and FDR<0.25) where input genes were ranked in descending order according to the Pearson’s correlation coefficient values derived between gene-level proteomic profile and *PI*_*RS*_.

Differential methylation probes located at promoter CGIs were identified by R package “*ChAMP*” ^34^, and probes that were hypermethylated (β>0.2) in any adjacent normal tissues were removed beforehand.

Individual fraction of copy number-altered genome (FGA), including gained (FGG) and lost (FGL) for TCGA-LIHC cohort was calculated as follows:

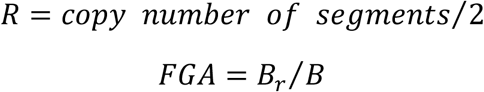

FGA is the genome fraction with log_2_(copy number)>0.3 versus the genome with copy number profiled; *B*_*r*_ represents the bases number in segments with absolute log_2_ *R* >0.3 and *B* denotes the bases number in all segments ^35^.

As previously described in our study ^36^, the R package “*RTN*” was used to reconstruct transcriptional regulons based on 71 cancerous chromatin remodelling regulators ^37^. A compendium of gene list consisted of 364 genes representing 24 microenvironment cell types was also retrieved from our previous study. Additionally, we collected 46 metabolism-relevant gene signatures from the literature ^38^. Gene set variation analysis (GSVA) was harnessed to quantify enrichment level for these gene sets through R package “*GSVA*”. The mRNA expression-based stemness index (mRNAsi) was calculated by a trained stemness index model based on one-class logistic regression machine-learning approach from single-sample perspective ^39^.

### Connectivity Map analysis and therapeutic response estimation

The Connectivity Map (CMap, https://clue.io/query) involves more than 7,000 gene expression profiles for approximate 1,300 compounds and assists discovering relationships between the diseases, cell physiology, and therapeutics ^40^. Therefore, potential therapeutic drugs for HBV-associated HCCs with high *PI*_*RS*_ were considered for those compounds with scores less than -95 (a negative score suggests a potential therapeutic effect). To validate the clinical implication, we employed R package “*pRRophetic*” to estimate the chemotherapeutic response for each HBV-associated HCC based on the drug sensitivity and phenotype data from PRISM as training cohort, and expression profiles of clinical tumors as testing cohort; AUC of each clinical tumor treated with specific compound was predicted by ridge regression, and 10-fold cross-validation was leveraged to evaluate predictive accuracy ^41^.

### Statistical analyses

All statistical analyses were conducted by R (Version: 4.0.2) using Fisher’s exact test for categorical data, Mann-Whitney U test for continuous data, and permutation test for the independence of two sets of variables measured on different scales. Correlation was measured via Pearson’s or Spearman’s coefficient under specified circumstances. Kaplan-Meier curves with log-rank tests were generated for survival rates comparison. Cox proportional hazard regression was used to estimate the hazard ratios (HRs) with 95% confidence intervals (95% CI). The prediction efficiency for 1-, 3- and 5-year survival was examined using receiver operating characteristic curve (ROC) by R package “*survivalROC*”. Model C-index was computed using R package “*survcomp*”. The restricted time survival (RMS) time ratio was estimated using R package “*survRM2*”. For all statistical analyses, a nominal *P<*0.05 was considered statistically significant.

## RESULTS

### Study design overview

A total of 606 HBV-associated HCC cases from five clinical cohorts were included. Of these patients, 536 patients have complete clinical follow-up. Detailed information was summarized in **Table S1**, and the entire study design was delineate in **Figure 1**.

**Figure 1.**
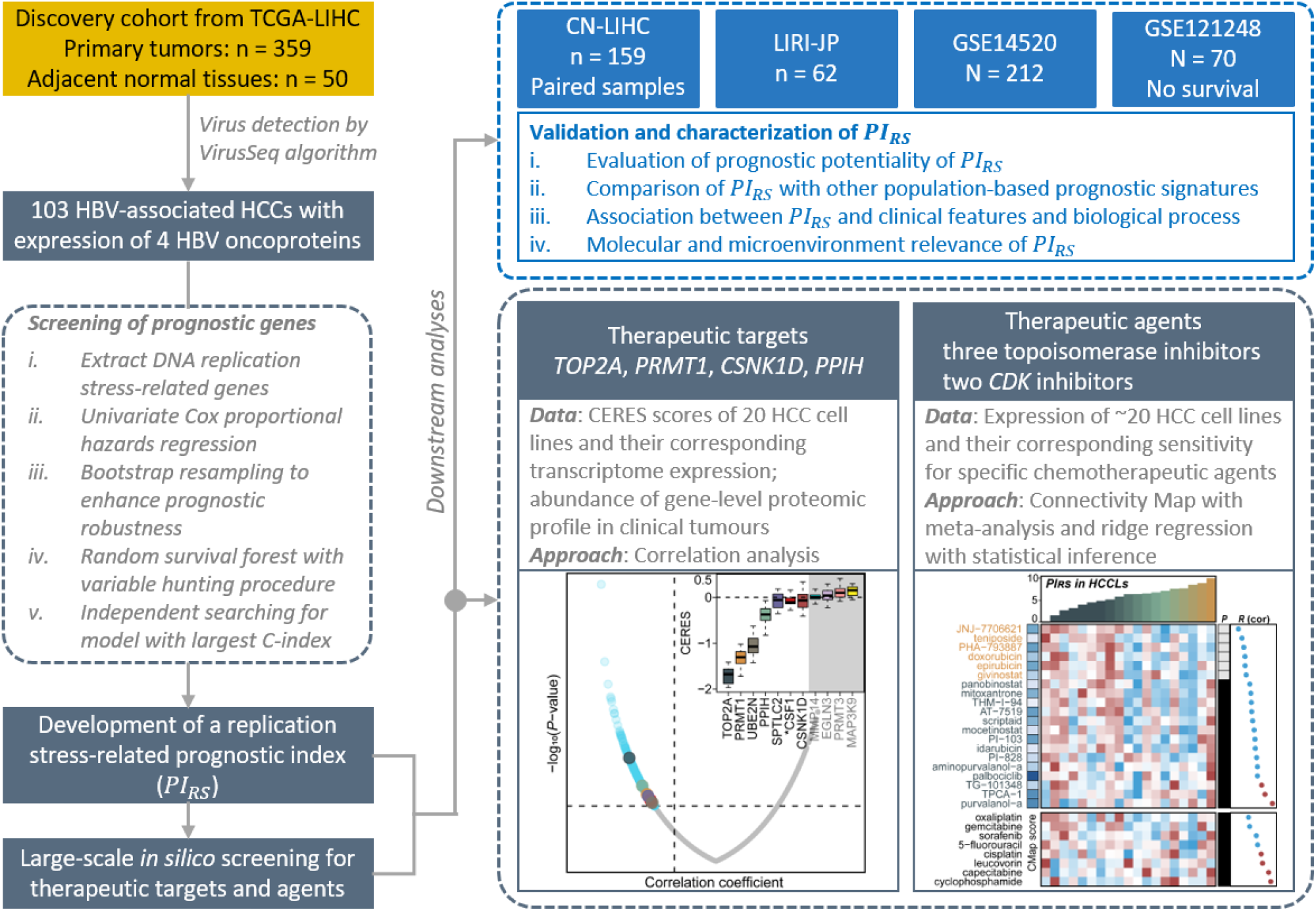
Design overview. This study enrolled a total of 606 HBV-associated HCC cases, and developed and validated a nine replication stress-related gene-based prognostic index (*PI*_*RS*_), which was further leveraged to predict potential therapeutic targets and agents.

### Development of nine replication stress-related gene-based *PI*_*RS*_ in HBV-associated HCCs

Based on genes within 21 replication stress signatures, we preliminarily identified 302 replication stress-related genes that were tightly associated with OS (*P*<0.01). To enhance prognostic robustness, bootstrap approach was conducted, resulting in 69 genes (*P*<0.01 in more than 80% resampling processes). Subsequently, RSF further narrowed down the list to a final panel of nine genes with the largest C-index (**Figure S2, Table S2**). A score of *PI*_*RS*_ was then calculated individually, ranging from 0 to 10. We developed the R package “*hccPIRS*” to calculate *PI*_*RS*_ from a single-sample perspective, which is documented and freely available at https://github.com/xlucpu/hccPIRS.

### Evaluation and validation of prognostic potentiality of *PI*_*RS*_

We reasoned that the area under the ROC curves were eligible (**Figure 2a-d**) given the time-dependent ROC at 1-, 3- and 5-year survival, the. Univariate and multivariate Cox regression were then conducted to assess the independent prognostic value of *PI*_*RS*_. Specifically, *PI*_*RS*_ combined with those significant clinical variables and frequently mutated genes (*i*.*e*., *TP53* and *CTNNB1*) in univariate analysis (*P*<0.05) were considered to construct a multivariate model (**Figure 2e**). In this manner, we found that *PI*_*RS*_ remained an independent prognostic factor after adjusting for other variables in TCGA-LIHC (HR: 1.25, 95% CI: 1.01-1.48, *P*=0.01), CN-LIHC (HR: 1.23, 95% CI: 1.07-1.43, *P*=0.005), GSE14520 cohorts (HR: 1.14, 95% CI: 1.02-1.27, *P*=0.024), but did not reach significance in LIRI-JP cohort which probably due to a relative small sample size (HR: 1.16, *P*>0.05). Additionally, *PI*_*RS*_ significantly stratified patients into low- and high-risk groups (*PI*_*LRS*_ and *PI*_*HRS*_) according to the up-tertile cut-off in TCGA-LIHC (HR: 5.07, 95% CI: 2.43-10.6, *P*<0.001; **Figure 2f, Figure S3a**), CN-LIHC (HR: 3.18, 95% CI: 1.87-5.38, *P*<0.001; **Figure 2g, Figure S3b**), LIRI-JP (HR: 3.96, 95% CI: 1.15-13.6, *P*=0.018; **Figure 2h, Figure S3c**) and GSE14520 (HR: 2.13, 95% CI: 1.38-3.3, *P*<0.001; **Figure 2i, Figure S3d**) cohorts; expression landscape of the nine genes was also validated in GSE121248 cohort (**Figure S3e**). The ratios of RMS ranging from 0.54 to 0.78 were observed in different cohorts (**Table S3**).

**Figure 2.**
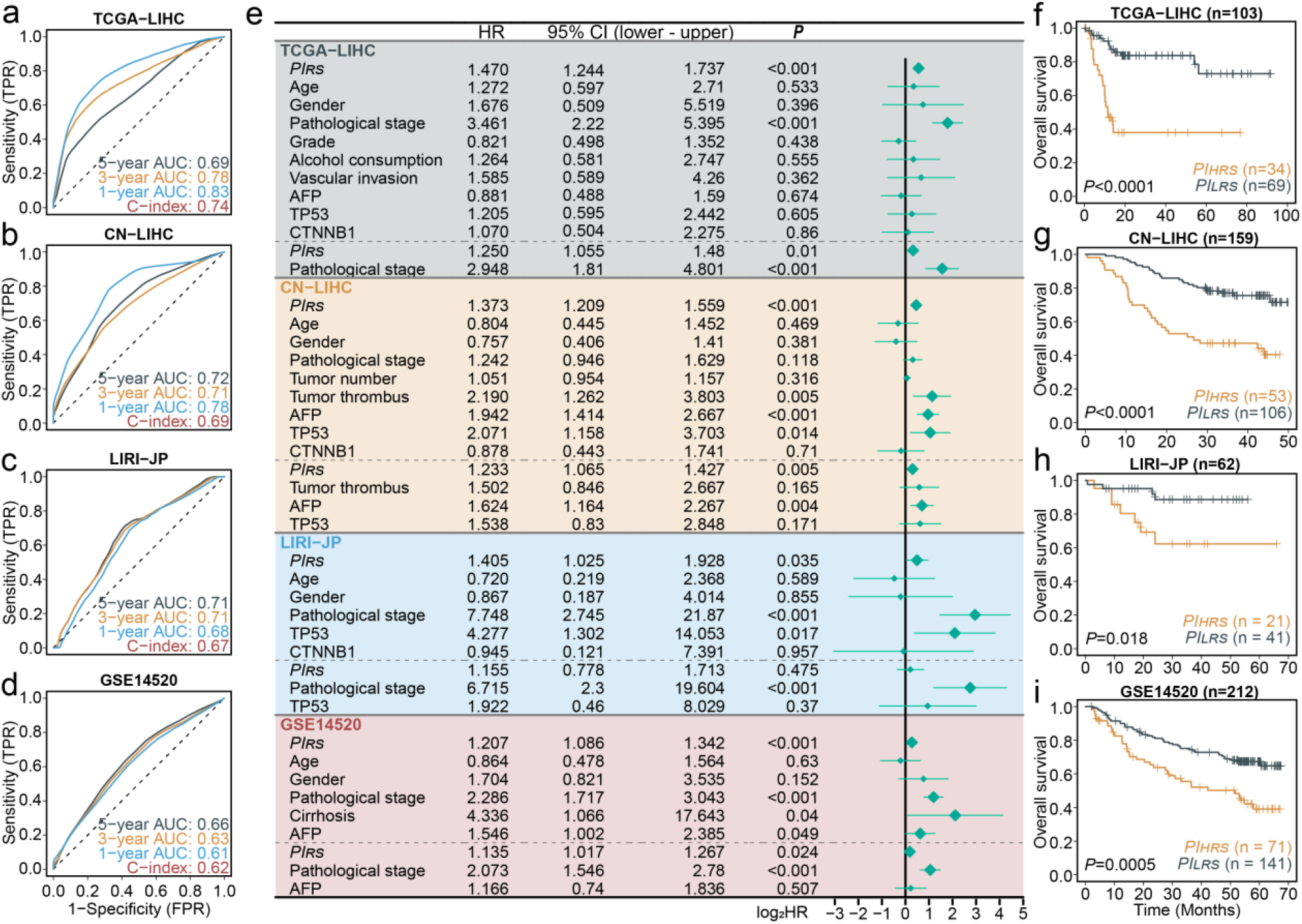
Performance of prognostic prediction based on *PI*_*RS*_ in four HBV-associated HCC cohorts. Time-dependent ROC curve analysis at 1-, 3-, and 5- survival for four different cohorts were shown in a) to d), respectively. e) Forestplot showing the hazard ratio (95% CI) in univariate Cox proportional hazards regression (above the dash line) and multivariate regression after adjusting for major clinicopathological features and the corresponding *P* values (below the dash line). Differentiate overall survival probability for patients stratified by *PI*_*RS*_ (*PI*_*HRS*_ and *PI*_*LRS*_ groups) was represented by Kaplan-Meier curves from f) to i) for TCGA-LIHC, CN-LIHC, LIRI-JP, and GSE14520 cohorts, respectively.

To test the universal prognostic value of *PI*_*RS*_, a general cut-off of 5.6 was determined using the up-tertile *PI*_*RS*_ among 606 HBV-associated HCCs. Using this cut-off, four cohorts were re-separated into *PI*_*LRS*_ * and *PI*_*HRS*_ * groups, and a strong association existed between the general cutoff-based new groups and cohort-specific cutoff-based groups (*P*<0.001; **Figure S4a**), implying the universal applicability of *PI*_*RS*_ in different cohorts/sequencing platform (**Figure S4b**). Consistently, the *PI*_*HRS*_* groups presented with significantly poorer OS than the matched *PI*_*LRS*_* groups (all, *P* < 0.05; **Figure S4c-f**).

We then compared *PI*_*RS*_ with other prognostic signatures in HCC. After 10,000 times of resampling, *PI*_*RS*_ showed comparable predictive performance as compared to other signatures in TCGA-LIHC cohort, while demonstrated comparable or superior performance in other HCC cohorts. Additionally, *PI*_*RS*_ maintained stable predictive performance across different cohorts (all, C-index>0.6), while other signature lost power in at least one cohort (**Figure S5, Table S4**).

### Association between *PI*_*RS*_ and replication stress signatures and chromatin remodeling regulons

We manifested that *PI*_*HRS*_ in TCGA-LIHC cohort showed significantly activated replication stress signatures at transcriptome-level (**Figure S6**). To further investigate transcriptomic differences, potential cancerous chromatin remodelling regulators were analyzed, reinforcing the biological pertinency of the up-tertile cutoff due to the remarkably shifted regulon activity pattern (**Figure S6**). Chromatin remodeling-associated regulon activity highlighted other possible differential regulatory schemas, suggesting that transcriptional networks driven by epigenome might be differentiators of great importance. Differential methylation analysis might further sustain the potential epigenetic differences between *PI*_*HRS*_ and *PI*_*LRS*_ because *PI*_*HRS*_ harbored more hypermethylated promoters than *PI*_*LRS*_ (2,867 *vs*. 136, FDR < 0.05; **Table S5**).

### Investigation of *PI*_*RS*_ associated clinical characteristics and biological processes

Considering that both TCGA-LIHC and CN-LIHC cohorts provide detailed clinicopathological information, we therefore investigated the association between *PI*_*RS*_ with clinical variables. Basically, higher *PI*_*RS*_ was tightly associated with higher alpha-fetoprotein (AFP) level and more aggressive clinical stage systems, including pathological stage and Barcelona Clinic Liver Cancer (BCLC) staging systems (**Figure 3a, Table S6-7**). Interestingly, we observed a mild and negative correlation between expression of oncoproteins HBVgp2-S (*R* = -0.24, *P*=0.015) and HBVgp3X (*R* = -0.28, *P*=0.005) with *PI*_*RS*_, as well as the *HBV*_*pca*_ in TCGA-LIHC cohort (*R* = -0.2, *P*=0.036; **Figure S7**).

**Figure 3.**
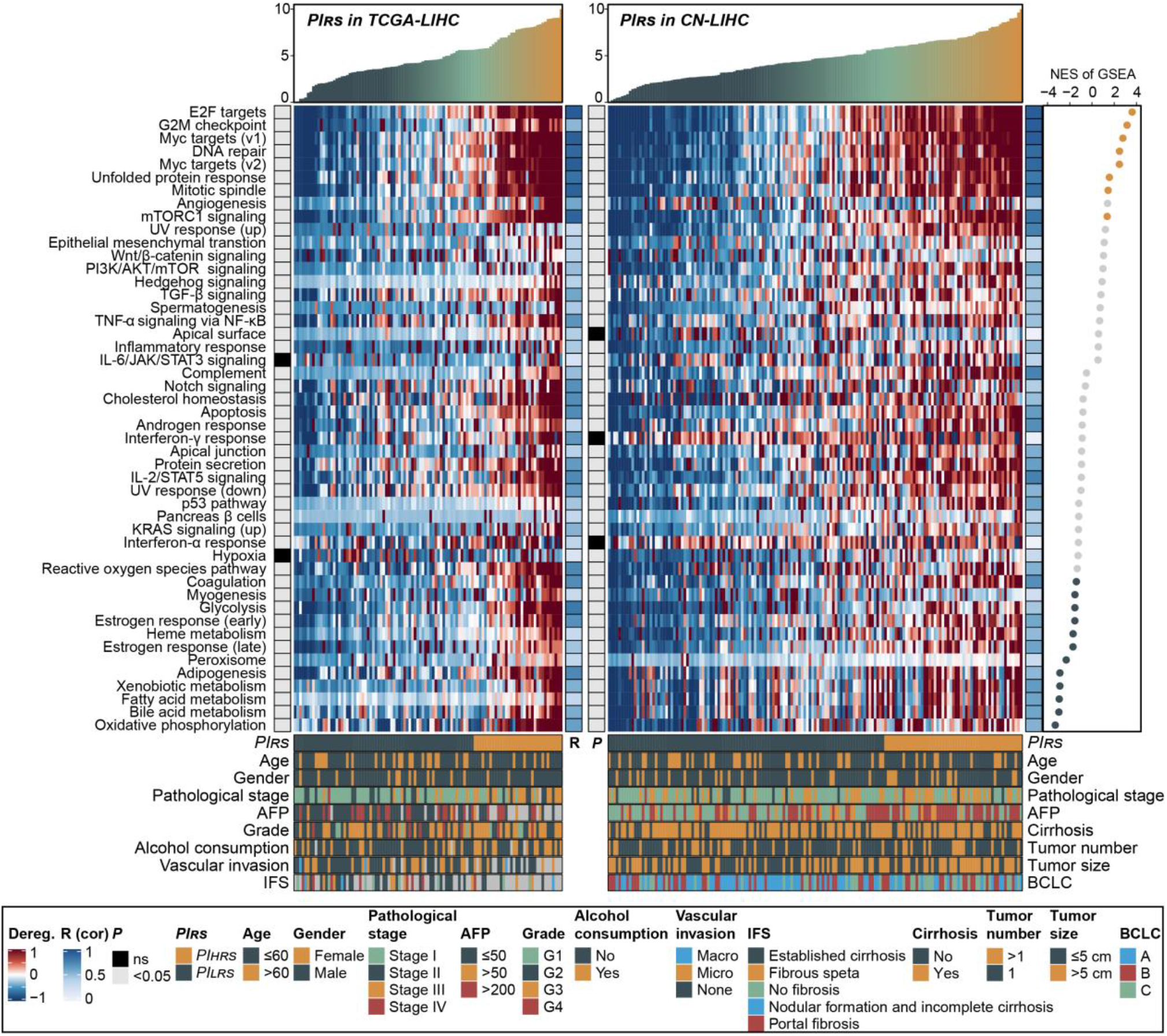
Landscape of *PI*_*RS*_ associated clinical characteristics and biological processes. The upper panel of the comprehensive heatmap shows the *PI*_*RS*_ in both TCGA-LIHC (left) and CN-LIHC (right) cohorts, and samples were arranged in an ascending sort. The middle panel of the heatmap displays the pathway deregulation scores of the cancer hallmarks that were quantified by “*pathifier*” algorithm at transcriptome level; the statistical *P* values and Pearson’s correlation coefficient between pathway deregulation scores and *PI*_*RS*_ were annotated on the left and right sides of the heatmap. The bottom panel presented the clinical annotation of each sample from each cohort. The right panel demonstrated to deregulation direction and its statistical significance through protein abundance-based GSEA upon the CN-LIHC cohort; yellow points for upregulated pathways, green points for downregulated pathways, and grey indicates non-significance. Dereg. stands for deregulation; NES stands for normalized enrichment score.

To investigate biological relevance of *PI*_*RS*_, we first quantified deregulation of cancer hallmarks through transcriptome-based Pathifier analysis based on TCGA-LIHC and CN-LICH cohorts considering their available normal samples. We found that almost all the hallmarks presented a significantly elevated degree of deregulation as *PI*_*RS*_ increased (**Figure 3**). Given that Pathifier did not specify the deregulation direction, we then performed gene-level protein abundance-based GSEA in CN-LIHC cohort; the results indicated that DDR-relevant and proliferation-related hallmarks were significantly upregulated as *PI*_*RS*_ increased, including G2-M checkpoint, *E2F* and *MYC* targets (**Figure 3**). Moreover, several metabolism-related processes were significantly activated as *PI*_*RS*_ decreased, including glycolysis and oxidative phosphorylation (**Figure 3**). To validate the *PI*_*RS*_-relevant biological process across different cohorts, we conducted a REM meta-analysis based on differential expression between *PI*_*HRS*_ and *PI*_*LRS*_ tumors in five cohorts (**Figure S8a**). We then performed transcriptome-based GSEA using pre-ranked gene list according to summary fold change, results of which verified the robustness of underlying biology of *PI*_*RS*_ (**Figure S8b**).

### Lower *PI*_*RS*_ tightly associated with activated immune/metabolism pathways and higher likelihood of responding to immunotherapy

Among 606 cases, Désert’s ECM/STEM and Boyault’s G3 subtype showed significantly higher *PI*_*RS*_ comparing other subtypes (both, *P*<0.001), which consistently mirrored the frequent *TP53* mutation as *PI*_*RS*_ increased (*P*<0.001; **Figure 4a-b**). Additionally, Désert’s periportal and Hoshida’s S3 subtype presented with significantly lower *PI*_*RS*_ as compared to other categories within the corresponding classification system (both, *P*<0.001; **Figure 4b**).

**Figure 4.**
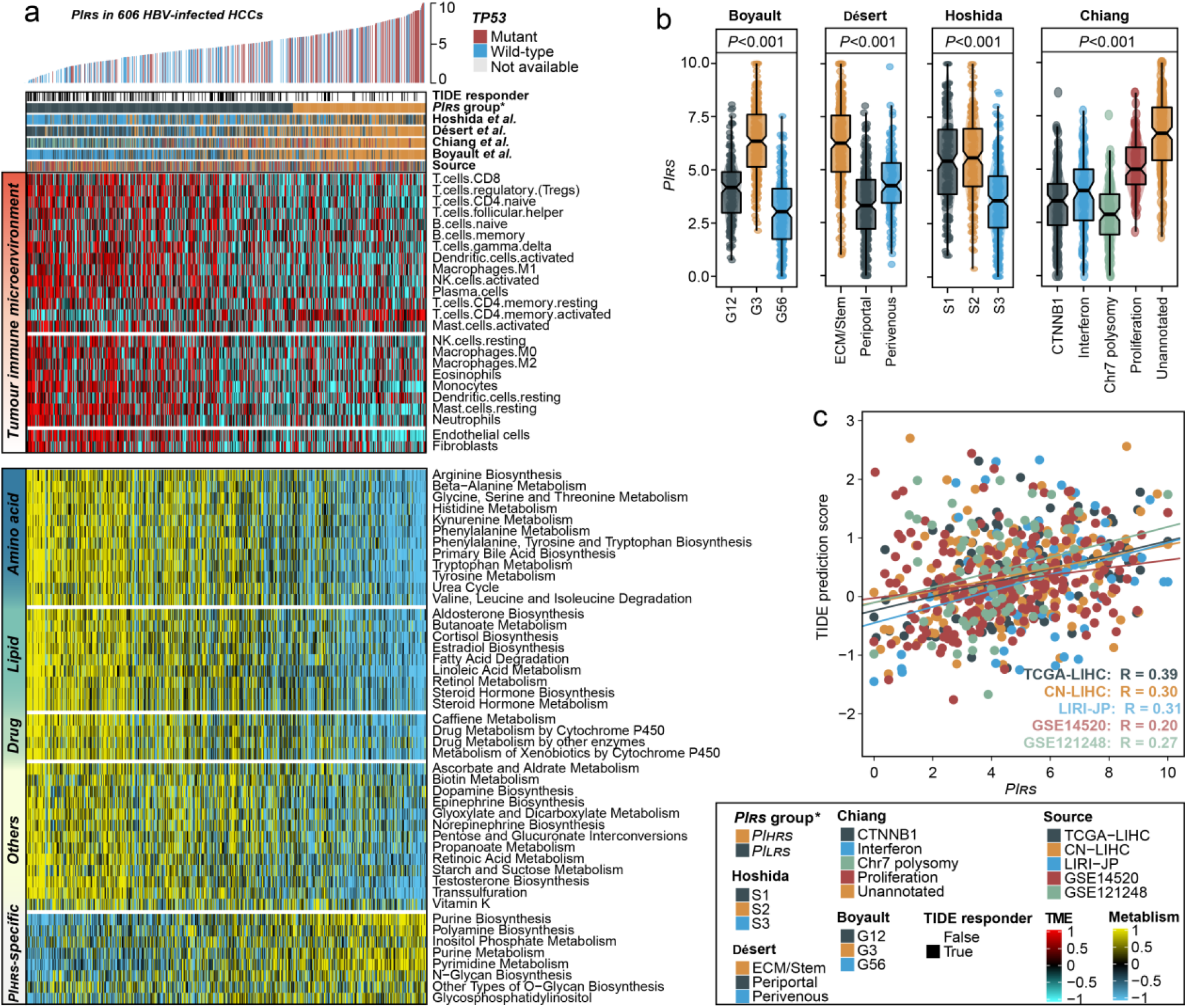
Association between immune/metabolism pathways, molecular features and *PI*_*RS*_. a) Heatmap showing the landscape of tumour immune microenvironment and metabolism pathways in 606 HBV-associated HCC cases from four independent cohorts. Samples were arranged in an ascending order according to the *PI*_*RS*_ and corresponding *TP53* mutation status was annotated; TIDE prediction and other previous molecular classification of HCC were annotated at the top of the heatmap. b) Boxplot showing the distribution of *PI*_*RS*_ in four molecular classification of HCC. c) Scatter plot showing the correlation between *PI*_*RS*_ and TIDE prediction score in four HBV-associated HCC cohorts; higher value of TIDE prediction score indicates less likely to benefit from anti-PD1/CTLA4 therapy.

Our previous study demonstrated that highly expressed human papillomavirus in cervical squamous cell carcinoma may stimulate the inflammatory/immune response of the host, leading to favorable prognosis ^20^; we therefore questioned whether this situation could be mirrored in HBV-associated HCC cases. We then quantified the infiltration level of 24 tumor microenvironment immune cells among 606 HCCs, and, strikingly, we found a globally activated immunocyte infiltration in *PI*_*LRS*_* group (**Figure 4a, Figure S9a**), which motivated us to investigate whether lower *PI*_*RS*_ was associated with higher likelihood of responding to immunotherapy. Considering that immune checkpoint inhibitors are not yet approved for HCC management by regulatory agencies, we therefore estimated the TIDE prediction score which represents the potential of tumor immune invasion (higher value indicates less likely to benefit from anti-PD1/CTLA4). We revealed that *PI*_*LRS*_ * group contained a significant higher proportion of TIDE-predicted responder than *PI*_*HRS*_* group (*P*<0.001; **Figure 4a**). Additionally, *PI*_*RS*_ showed significant and positive correlation with TIDE prediction score in all five HCC cohorts, (**Figure 4c**), suggesting that HBV-associated HCC patients with lower *PI*_*RS*_ could possibly respond to immune checkpoint inhibitors.

Additionally, the cancerous genomic landscape has been manifested being related to anti-tumour immunity; for instance, presence of neoantigen triggers T-cell responses ^42^, whereas aneuploidy may cause immune evasion and attenuation of immunotherapy response ^43^. In this context, we investigated TCGA-LIHC cohort and found *PI*_*RS*_ showed strong and negative correlation with the number of predicted neoantigen (*R*=-0.38, *P*=0.027; **Figure S9b**) while positively correlated with number of broad-level deletion (*R*=0.44, *P*<0.001; **Figure S9c**) and FGL (*R*=0.28, *P*=0.005; **Figure S9d**); no statistical significance was obervsed regarding FGG or broad-level amplification (both, *P*>0.25, not shown).

Considering the inducement of the non-infiltrated phenotype in *PI*_*HRS*_* group, a recent study reported that the transcription factor nuclear factor erythroid 2-related factor 2 (*NRF2*) contributes to an “immune-cold” phenotype by inducing *COX2*/*PGE2* and inhibiting the DNA-sensing innate immune response ^44^; *NRF2* also accumulated in the nucleus and formed foci at DNA damage sites, thus facilitating DDR and DNA repair ^45^. Therefore, we investigated the expression of *NRF2* and *COX2* of three cohorts (*i*.*e*., TCGA-LIHC, CN-LIHC, and GSE121248) in which both genes are matchable. We found that *PI*_*HRS*_ groups showed significantly higher expression of *NRF2* and its downstream marker *COX2* in TCGA-LIHC (*P*=0.009 for *NRF2, P*=0.001 for *COX2*; **Figure S9e**) and CN-LIHC cohorts (*P*<0.001 for *NRF2, P*<0.001 for *COX2*; **Figure S9f**); we did not observe a statistical significance of *NRF2* in GSE121248 cohort (*P*=0.15, not shown), *COX2* was dramatically upregulated in *PI*_*HRS*_ group (*P*=0.042; **Figure S9g**), though.

As we have already shown that metabolic pathways could be suppressed with increasing *PI*_*RS*_, we then investigated the metabolic landscape among 606 HCCs. As expected, patients belonging to *PI*_*LRS*_* presented with significant activation of metabolism relevant signatures, including amino acid metabolism relevant signatures, lipid metabolism relevant signatures, and drug metabolism relevant signatures; while only several metabolic pathways were enriched in *PI*_*HRS*_* group (**Figure 4a, Figure S9h**). The activation of metabolic pathways and enrichment of periportal HCC subtype indicated that HCC cases in *PI*_*LRS*_* group preserve the default metabolic program (*e*.*g*., gluconeogenesis, amino acid catabolism, and urea cycle) of normal liver and are well-differentiate and non-proliferative, which may synergistically contribute to a favorable prognosis.

We verified the non-proliferative nature of tumors with lower *PI*_*RS*_ due to their significant lower expression of the proliferation marker—*Ki-67* (all, *P*<0.001; **Figure S10a**). Additionally, we found that *PI*_*RS*_ showed strongly positive correlation with mRNAsi scores (*R*=0.45, *P*<0.001; **Figure S10b-c**), which indicated patients with high *PI*_*RS*_ might resistant to conventional chemotherapy and radiation therapy ^39^, thereby emphasizing the essentiality of tailoring therapeutic strategies for patients with high risk.

### Identification of potential drug targets for HBV-associated HCCs with high *PI*_*RS*_

Proteins that are strongly positively correlated with high *PI*_*RS*_ may hold potential therapeutic implications for a subset of HBV-associated HCCs under high mortality risk. Unfortunately, most human proteins currently lack active binding sites for small molecule compounds, which impedes them from becoming potent drug targets. Thus, using the target information of thousands of compounds ^25^, a total of 1,382 drug targets corresponding to 6,706 chemical compounds were screened through a two-step analysis (**Table S8**). Specifically, we first conducted Pearson’s correlation between the transcriptome expression of target (gene) and *PI*_*RS*_ in TCGA-LIHC and CN-LIHC cohorts, respectively. Totally 369 common genes with correlation coefficient >0.3 (*P*<0.05) was considered as *PI*_*RS*_-related drug targets (**Figure 5a, Table S9**). Subsequently, *PI*_*RS*_ was further calculated for each HCCL, and Spearman’s correlation analysis was performed between CERES and *PI*_*RS*_ of HCCLs (**Figure 5b, Table S10**). Given that a lower CERES score indicates a higher likelihood that the gene is essential in cell growth and survival, a *PI*_*RS*_-related gene was considered as disease progression-dependent drug target if its correlation coefficient <-0.3 (*P*<0.05). In this manner, 11 genes were identified, containing *TOP2A, PRMT1, EGLN3, CSF1, CSNK1D, MMP14, MAP3K9, PPIH, SPTLC2, UBE2N*, and *PRMT3*. Of note, genes of *EGLN3, MMP14, MAP3K9*, and *PRMT3* had averaged CERES scores greater than zero (**Figure 5b**), suggesting their less fundamental nature in HCC development. We further investigated the remaining seven genes for their protein abundance using CN-LIHC cohort, and *TOP2A* (*R*=0.78), *PRMT1* (*R*=0.45), *CSNK1D* (*R*=0.23), and *PPIH* (*R*=0.35) might be potentially druggable considering their significant and positive correlation with *PI*_*RS*_ (all, *P*<0.05; **Figure 5c**). The strong and positive correlation between these potential gene targets and corresponding *PI*_*RS*_ in other HBV-associated cohorts were also validated (**Figure 5d-h**).

**Figure 5.**
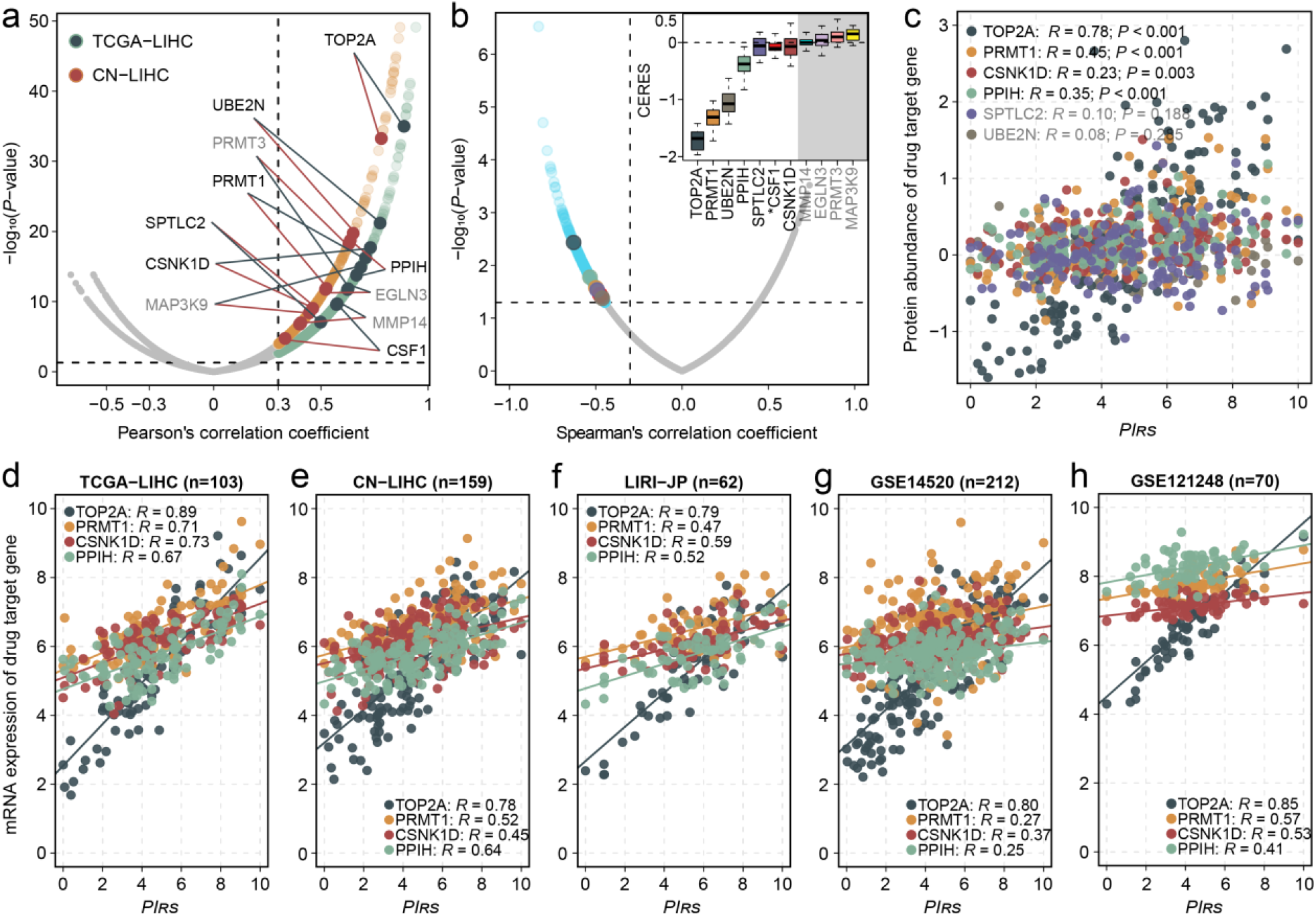
Identification of *PI*_*RS*_ -related therapeutic targets. a) Volcano plot showing the correlation coefficient against statistical significance derived from Pearson’s correlation analysis between *PI*_*RS*_ and therapeutic target expression at transcriptome-level in clinical tumors. Light-colored points represent potential targets that pass the threshold (*R*>0.3 and *P*<0.05), and dark-colored points represent targets that were also identified from CERES analysis; target name in grey color indicates potential disease progression independency. b) Volcano plot showing the correlation coefficient against significance derived from Spearman’s correlation between *PI*_*RS*_ and CERES scores of drug targets in HCC cell lines. Light blue points represent potential targets that pass the threshold (*R*<-0.3 and *P*<0.05), and colored points are those shared with former correlation analysis in clinical samples. Distribution of CERES scores of identified target among HCC cell lines were positioned as boxplot at the top right corner; targets with averaged CERES scores of greater than 0 were colored in grey, and target marked with an asterisk indicated unmatchable in gene-level proteomic profile. c) Scatter plot showing the association between *PI*_*RS*_ and gene-level protein abundance of drug target in CN-LIHC cohort. Scatter plots of the association between *PI*_*RS*_ and target expression at transcriptome-level for five cohorts were shown in d) to h), respectively.

### Therapeutic response prediction of targeted chemotherapy

To screen potentially effective chemical compounds to HBV-associated HCCs with high *PI*_*RS*_, we performed CMap analysis as a preliminary to investigate the therapeutic potential of candidate agents. To this end, 150 up-regulated and 150 down-regulated genes with the most significant fold changes in REM meta-analysis were submitted to CMap, which identified 85 chemical compounds under pharmacologic CMap classes (**Table S11**); a total of 30 agents showed perturbagens with enrichment scores below -95, including topoisomerase inhibitors, *CDK* inhibitors, *HDAC* inhibitors, *PI3K* inhibitors, *etc* (**Figure 6a**).

**Figure 6.**
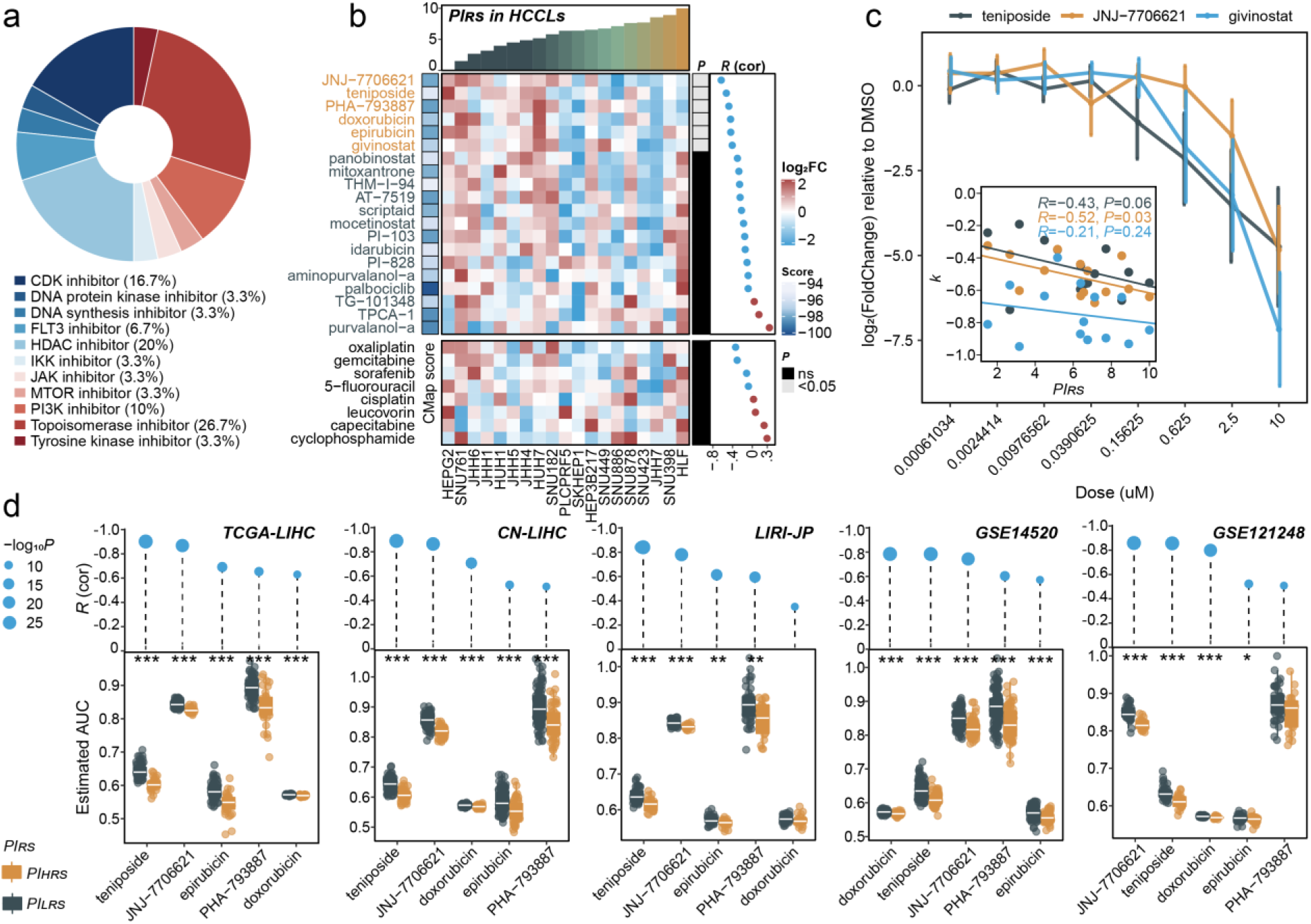
Identification of candidate therapeutic agents with higher sensitivity in patients with high *PI*_*RS*_. a) Pie chart showing the fraction of chemical compounds according to the under pharmacologic CMap classes. b) Heatmap showing the distribution of drug response in HCC cell lines ordered by *PI*_*RS*_ in an ascending sort. The upper heatmap represents the compounds identified by CMap with scores less than -95 and the top six drugs are marked in yellow due to the statistical significance of correlation between drug sensitivity and *PI*_*RS*_, and others were marked in blue; the bottom heatmap shows the drugs that commonly used for treating HCC. CMap scores were annotated at left, Spearman’s correlation coefficient and statistical *P* values were annotated at right. c) Broken line chart showing the drug sensitivity at eight dose points; each broken point was presented as mean ± standard deviation. The association between the slope *k* (linear regression between dose and drug response) and *PI*_*RS*_ were shown as scatter plot at the bottom left corner. d) Boxplot showing the distribution of inferred sensitivity for five candidate therapeutic agents (represented by AUC) between *PI*_*HRS*_ and *PI*_*LRS*_ groups in five HBV-associated HCC cohorts; coefficient and significance of Pearson’s correlation between predicted AUC and *PI*_*RS*_ were annotated at the top of boxplot.

In order to enhance the credibility of these drug inferences, we searched these candidate agents in DepMap, leading to 20 matched compounds. Next, we calculated the Spearman’s correlation coefficient between primary drug response (measured as log_2_FoldChange) and *PI*_*RS*_ in 19 HCCLs, which yielded six drugs whose sensitivity significantly elevated with the increase of *PI*_*RS*_ (all, *R*<-0.4, *P*<0.05; **Figure 6b**), including JNJ-7706621, teniposide, PHA-793887, doxorubicin, epirubicin, and givinostat. Of note, teniposide, a Topoisomerase inhibitor, targets *TOP2A* which we have already considered potentially druggable, suggesting that our *in silico* strategy for drug screening could be reliable. Additionally, we investigated the association between *PI*_*RS*_ of HCCLs and drug sensitivity of the most common chemotherapy for treating HCC, containing sorafenib, gemcitabine, oxaliplatin, cisplatin, 5-fluorouracil, capecitabine, leucovorin, and cyclophosphamide ^46^; while none of them showed significant correlation with *PI*_*RS*_ (all, *P*>0.05; **Figure 6b**), implying that these routine interventions might fail to pose a prognosis-dependent effect in HBV-associated HCC patients.

Notably, we also matched 3 out of 6 drugs (*i*.*e*., teniposide, JNJ-7706621, and givinostat) with second measurement for dose-finding in 14 HCCLs; as expected, as the dose increased, the responsiveness of HCCLs to the specific drug elevated (**Figure 6c**). Additionally, we found that the dose-dependent sensitivity (measured as slope *k* of linear regression) of teniposide (*R*=-0.43, *P*=0.06) and JNJ-7706621 (*R*=-0.52, *P*=0.03) is dramatically associated with *PI*_*RS*_ in a negative manner, suggesting that high *PI*_*RS*_ might synergistically promote the antitumor activity of these compounds (**Figure 6c**).

We further validated the clinical implication of these 5 drugs (removal of givinostat) through a model-based strategy. Generally, in addition to the observation where correlation analysis between *PI*_*RS*_ and predicted AUC demonstrated widely negative association in HBV-associated HCCs, patients belonging to *PI*_*HRS*_ groups showed remarkably lower estimated AUC than those in *PI*_*LRS*_ groups (**Figure 6d**).

## DISCUSSION

HCC, especially HBV-associated, remains the leading cause of cancer-related mortality worldwide. The genome instability caused by HBV integration might trigger malignancies to present chronic replication stress, thereby providing an exploitable therapeutic vulnerability for HCC ^47^. We herein aimed to develop an efficient approach of prognostic prediction for HBV-associated HCC based on DNA replication stress signatures, and investigate tailored therapeutic strategy for patients with high mortality risk, which is of great significance to maximize benefit from precise medicine. In this context, we proposed *PI*_*RS*_, a machine-learning trained prognostic predictor for HBV-associated HCC. Apart from being informative regarding prognosis, *PI*_*RS*_ can be also leveraged for precise oncology, as a biomarker to guide targeted treatment. Specifically, four potential therapeutic targets (*i*.*e*., *TOP2A, PRMT1, CSNK1D*, and *PPIH*) and five agents including three topoisomerase inhibitors (*i*.*e*., teniposide, doxorubicin, and epirubicin) and two *CDK* inhibitors (*i*.*e*., JNJ-7706621 and PHA-793887) were identified for patients with high *PI*_*RS*_.

Type II topoisomerases (TOP2) are pervasive enzymes that can alter DNA superhelicity and unlink replicating DNA ^48^. In HCC, Panvichian *et al*. revealed that *TOP2A* is significantly overexpressed in tumor tissues as compared to adjacent normal tissue, and another meaningful result is that high *TOP2A* expression is more observed in patients with the positive serum HBsAg test ^49^. The protein arginine methyltransferase 1 (*PRMT1*) may play a pivotal role in multiple cellular processes, including proliferation, transformation, invasiveness, and survival of malignancies through methylation of arginine residues that underlie these processes ^50^. *PRMT1* can promote the tumorigenesis and progression of HCC through activating *STAT3, TGF-β1*/*Smad* and *HNF4α* pathways ^51-53^. Moreover, it has recently been reported that genetic knockdown and pharmacological inhibition of *PRMT1* by DCPT1061, a novel potent inhibitor, drastically induced G1-phase cell cycle arrest and suppressed cell growth of clear cell renal cell carcinoma ^54^; another *PRMT1* inhibitor (GSK3368715) was capable of impairing replication restart of pancreatic ductal adenocarcinoma, thus inhibiting tumor growth ^55^. The casein kinase 1 delta (*CK1δ, CSNK1D*) is a member of serine/threonine protein kinase family that comprises of six isoforms (*i*.*e*., α, δ, ε, γ1, γ2 and γ3) that were involved in several signaling pathways (*e*.*g*., Hedgehog, Wnt, and Hippo), and mediates numerous cellular processes (*e*.*g*., DNA replication, DDR, RNA metabolism, membrane trafficking, cytoskeleton maintenance, and circadian rhythm) ^56^. Rosenberg *et al*. revealed that silencing or inhibition of *CSNK1D* using SR-3029 provokes potent anti-tumor effects for breast cancer cells *in vivo* ^*57*^. As to *PPIH*, the protein encoded by which is a member of the peptidyl-prolyl cis-trans isomerase (PPIase) family. Although limited evidence directly demonstrated anti-tumor effect of inhibiting *PPIH*, Uchida *et al*. revealed inhabitation of *PIN1*, the popular member of PPIase family impaired the growth of several cancer cell lines ^58^. Taken together, emerging evidences have shown that the four targets we identified all play special roles in malignant development and several inhibitors have demonstrated potent anti-tumor effect in specific cancer type, suggesting the feasibility of developing corresponding targeted therapies for high-risk HBV-associated HCC.

The inhibition of *TOP2* (topoisomerase inhibitor) is a therapeutic strategy for cancer treatment and have been applied to treat cancers for many years, such as etoposide and teniposide that targets *TOP2A*. Several clinical trials are ongoing to evaluate the effectiveness of topoisomerase inhibitors in HCC (*e*.*g*., NCT00351195, NCT03533582, NCT03017326). Doxorubicin and epirubicin are traditional topoisomerase inhibitors, and has been widely used for the treatment of HCC. Even though they have never been recommended for systemic treatment of HCC, these two remain the main drugs used for transarterial chemoembolization (TACE) of HCC ^59, 60^. Unfortunately, a recent result from the phase III Alliance/CALGB 80802 trial failed to observe the benefit from the addition of doxorubicin treatment to sorafenib, and the safety concerns of doxorubicin raised the cautious attitude for its limited application, due to the presence of underlying cirrhosis, hematologic toxicity, and cardiac toxic events ^61^; such disappointing report inversely lays more emphasis on the desire of developing robust biomarkers for precision medicine, thus enhancing efficacy and reducing adverse reactions. Based on our findings, we revealed that HBV-associated HCC patients with high *PI*_*RS*_ are more susceptible to topoisomerase inhibitors (*i*.*e*., teniposide, doxorubicin and epirubicin), which might guide relevant clinical trial designs in the future.

JNJ-7706621, a potent *CDK* inhibitor targeting *CDK1*/*2*, blocks tumor progression through cell cycle, causing cells to accumulate in G2/M phase, preventing cells from entering mitosis and activating apoptosis, which could be useful for treating various cancers ^62, 63^. PHA-793887 is another kind of multiple *CDK* inhibitor with the activity against of *CDK1*/*2*/*4*/*5*/*7*/*9*, which is also revealed in our study as a potential drug for high-risk HBV-associated HCCs. Brasca *et al*. demonstrated that PHA-793887 had good efficacy in the human ovarian A2780, colon HCT-116 and pancreatic BX-PC3 xenograft models and was well-tolerated via daily intravenous treatment, suggesting PHA-793887 is promising as a drug candidate for clinical evaluation ^64^. In fact, the inhibitors of *CDK* pathway have been widely reported with the function of inducing apoptosis and ongoing test in the clinical trials for HCC, including palbociclib (NCT01356628, *CDK4*/*CDK6* inhibitor), flavopiridol (NCT00087282, *CDK1*/*2*/*4*/*6*/*7* inhibitor) and milciclib (NCT03109886, *CDK2*/*4*/*5*/*7* inhibitor). Additionally, Ehrlich *et al*. revealed that the combinational value of *CDK5* inhibitor, roscovitine, with DNA-damage-inducing chemotherapeutics synergistically inhibited HCC tumor progression ^65^. Tourneau *et al*. also observed a partial response of selicicib, a *CDK1*/*2*/*7*/*9* inhibitor, in one HCC patient from phase I evaluation ^66^. Therefore, we reasoned that the pan-*CDK* inhibitor PHA-793887 might achieve better response for HBV-associated HCC cases with high *PI*_*RS*_.

We acknowledged several limitations. Foremost, the enrolled cohorts varied in size, composition, and sequencing technology. Besides, incomplete treatment records may bias the study design. Ultimately, bulk sequencing and microarray profiles are confounded by signals quantified from a mixed cell populations and are distinct from cell lines concerning expression and drug sensitivity; thus, combining our findings with multiplex immunohistochemistry to delve intrinsic cancer cell alterations and their crosstalk with the tumour microenvironment that dictate therapeutic response is warranted.

## CONCLUSIONS

In summary, we developed *PI*_*RS*_, a novel DNA replication stress signature, which serves as a single-sample survival predictor for HBV-associated HCC and may be readily translated to clinical practice to guide prognosis stratification and personalized therapeutic strategies. Specifically, physicians could adopt a low-toxicity therapeutic strategy to avert overtreatment for low *PI*_*RS*_ patients who may also benefit from immunotherapies or metabolic therapies. To those patients with high *PI*_*RS*_, our study identified potential therapeutic targets and agents, which might improve their clinical outcomes in a more effective way. Overall, current work has not only shed new insight to prognostic stratification, but also provided a roadmap for the rational clinical development of personalized treatment.

## KEY POINTS

- This study highlights the genetic heterogeneity of HBV-associated hepatocellular carcinoma concerning prognostic DNA replication stress.
- This study developed a tailored prognostic index that could improve the population-based prognostication approach.
- This study manifested that a prognostic index enables exploitable therapeutic vulnerabilities for patients with high mortality risk.
- This study identified four therapeutic targets and five agents (three topoisomerase inhibitors and two *CDK* inhibitors) for HBV-associated hepatocellular carcinoma.

## DECLARATIONS

### Ethics approval and consent to participate

As the data used in this study are publicly available, no ethical approval was required.

### Consent for publication

Not applicable.

### Availability of data and material

The raw data for this study were generated at the corresponding archives and were detailed for source in the section of *Materials and Methods*. We developed the R package, “*hccPIRS*”, which is documented and freely available at https://github.com/xlucpu/hccPIRS. This package calculates replication stress-related prognostic index (*PI*_*RS*_) for HBV-associated HCC patients, and estimates the enrichment of 21 replication stress signatures. If specified, a heatmap will be generated to show the landscape of the replication stress signatures in an ascending order of *PI*_*RS*_ score. Core R codes of prognostic model development, functional enrichment, REM-meta analysis, and drug target identification have been uploaded to GitHub (https://github.com/xlucpu/hccPIRS/tree/main/code). Derived data and other analytic code supporting the findings are available from the corresponding author [F. Y] on reasonable request.

## Authors’ contributions

Conceptualization, X. L, J. M and Y. Z; methodology, X. L, H. W, and L. J; formal analysis, X. L, J. M, Y. Z, H. W, X. R, Y. C, Q. Y, and L. S; investigation, all authors; writing the original draft, X. L, J. M and Y. Z; visualization, X. L; funding acquisition, F. Y, and supervision, L. J, and F. Y.

## Acknowledgements

We greatly appreciate the patients and investigators who participated in the corresponding medical project for providing data.

